# Parasitic-plant parasite utilizes flowering pathways at unconventional stages to form stem-derived galls

**DOI:** 10.1101/2024.10.17.618901

**Authors:** Naga Jyothi Udandarao, Yuki Yamashita, Ryo Ushima, Tsutomu Tsuchida, Kanako Bessho-Uehara

**Affiliations:** Graduate School of Life Sciences, Tohoku University, Sendai, Miyagi 980-8578, Japan; Graduate School of Science and Engineering for Education, University of Toyama, 3190 Gofuku, Toyama City 930-8555, Toyama, Japan; Faculty of Science, Academic Assembly, University of Toyama, 3190 Gofuku, Toyama City 930-8555, Toyama, Japan

**Keywords:** Gall, *Cuscuta campestris*, *Smicronyx madaranus*, cell cycle, flowering, photosynthesis

## Abstract

Galls induced by various organisms exhibit diverse morphological and physiological characteristics, involving complex plant-insect interactions. Most transcriptome analyses to date have focused on leaf-derived galls. To better understand gall formation mechanisms, we investigated stem-derived galls induced by the weevil *Smicronyx madaranus* on the parasitic plant *Cuscuta campestris* at gene expression, cellular, and physiological levels. RNA-seq across four developmental stages identified differentially expressed genes and associated gene ontology terms. Consistent with histological observations, genes related to cell division and the cell cycle were upregulated early but decreased as the gall matured. Similar to leaf-derived galls, we found high expression of *PLETHORA* and meristem-related homeobox genes in early gall development, suggesting that stem cell induction and maintenance are involved in various gall types. Like leaf-derived galls, the expression of genes related to floral organ development increased through the gall development. However, their expression patterns were dramatically different: downstream genes in the flowering pathway were highly expressed at the initial gall stage, whereas upstream genes were highly expressed later. This suggests that the weevil might activate the flowering pathway at unconventional stages, potentially rerouting the typical flowering cascade to influence gall development. Unlike the decrease in photosynthesis-related genes in leaf-derived galls, we observed an increase in these genes in galls formed on the stem of the holoparasitic plant. Shading experiments confirmed that photosynthesis is crucial for both gall growth and the weevil. This study highlights how gall-inducers can co-opt host resources and genetic pathways, offering new insights into the complexity of plant-insect interactions.

## Introduction

Galls are abnormal plant outgrowths induced by various organisms, including bacteria, fungi, mites, and insects. Insects are particularly noteworthy for their diversity and prevalence in gall formation, with approximately 211,000 species capable of inducing galls under various environmental and host conditions (Espírito-Santo and Fernandes, 2007; Desnitskiy et al., 2023). These structures can develop on different plant organs, including leaves, stems, roots, and flowers, reflecting the complex relationship between insects and host plants (de Oliveira et al., 2014). Galls provide a unique niche that offers shelter and nutrition to the gall-inducing insects. They create a controlled microenvironment, protecting the larvae from external environmental stresses and predators (Harris and Pitzschke, 2020).

The formation of galls is a highly organized process, typically involving four developmental stages: initiation, growth, maturation, and senescence (Rohfritsch and Shorthouse, 1982). It is intriguing how insects manipulate each stage along with responding to plant developmental processes resulting in complex gall structures. Galls typically consist of three tissue types: stem cells, vascular tissues, and protective tissues (Stone and Schönrogge, 2003; Giron et al., 2016). During gall formation, meristematic stem cells are controlled in their proliferation and differentiation to generate the specific tissues that compose the gall. For instance, phylloxera utilizes undifferentiated stem cells associated with leaf veins to develop galls (Schultz et al., 2019). Meristematic marker genes such as class-1 *KNOTTED-like homeobox* (*KNOX*) transcription factors are often activated in the early stages of gall development, as seen in *Rhus javanica* galls induced by the aphid *Schlechtendalia chinensis* (Hirano et al., 2020). Interestingly, flowering-related genes are upregulated in the initial stage of galls across distant plant species (Takeda et al., 2021). Auxin, cytokinin, and other hormone-responsive genes likely collaborate with these transcription factors to drive complex morphological features in galls induced by insects (Bartlett and Connor, 2014; Body et al., 2019).

However, most transcriptome analyses have been restricted to leaf-derived galls, leaving stem-derived galls underexplored. To characterize the differences based on the derived plant organ, it is essential to obtain precise transcriptome data from stem-derived galls. Furthermore, gall development in natural field conditions presents significant challenges for analyzing developmental stages or gene expression patterns, complicating the control of variables that influence gall formation. Thus, model systems for gall research are necessary to explore the detailed molecular mechanisms.

In a previous study, we established a new model system consisting of the parasitic plant *Cuscuta campestris* and the stem-galling weevil *Smicronyx madaranus* (Murakami et al., 2021). This system allows year-round stable cultivation in the laboratory. In this model, gall induction occurs within a few days, and the galls enlarge and fully develop into fruit-like structures within two weeks. RNA interference (RNAi) has proven effective for gene function analysis in *S. madaranus* (Ushima et al., 2024), enabling detailed molecular analysis of gall formation. Observation of the development of *C. campestris* galls indicated that the photosynthetic activity increases within the galls, leading to the accumulation of starch, which serves as food for the larvae (Murakami et al., 2021). This is in contrast to previous findings in leaf-derived galls, where photosynthesis-related genes are typically suppressed, converting leaf tissues from nutrient sources to sinks (Takeda et al., 2019; Hirano et al. 2020). The reverse process observed in *C. campestris* stem-galling system offers a unique opportunity to explore the diversity of gall formation mechanisms.

In this study, we aimed to investigate the molecular mechanisms underlying gall formation in *C. campestris*, focusing on gene expression and physiological aspects of this plant, with a particular emphasis on the flowering pathway. The gall-inducing system in the lab has enabled time-course histological analyses and experimental manipulations, allowing us to understand gall development deeply. Unlike previous studies that have mostly focused on early-stage gene expression patterns, this system’s advantages facilitated the examination of not only the early stages but also the middle and late stages of gall development. This approach revealed developmental similarities at the cellular level akin to fruit development. Additionally, we examined the extent to which photosynthesis in the gall affects gall and larval development.

## Results

### 3D structure of different stages of *C. campestris* gall

Galls are induced at the nodes of young stems of *C. campestris* when the mother of *S. madaranus* lays eggs, causing the galls to swell over time (**Fig. 1A-D, Supplementary** Fig. 1A). During the Sgall stage, three elongated spaces were found in the center of the gall by X-ray micro-computed tomography (micro-CT) observation (**Fig. 1E**). Near the eggs, an object with X-ray absorption similar to that of the egg was always located on the epidermal side, implying that this lid-like structure may have been created by the mother and functioned as a cover for the oviposition hole (**Fig. 1F**). The oviposition hole was observed at Sgall stage (**Supplementary Fig. S1B**), but it was difficult to observe at the Mgall stage due to repair or cell enlargement as the gall grows (**Supplementary Fig. S1C**). One to several larvae were observed per gall, and larvae were present inside of larval chamber (**Fig. 1G**). Several egg- or larval-free holes were observed at all gall stages. The size of these holes increased as the stage progressed. This indicated that rapid tissue enlargement may create spaces where the supply of cells cannot keep up (**Fig. 1H**). In the Lgall stage, larvae within the chambers had significantly developed (**Fig. 1I**). Additionally, the number of vascular bundles originally present in the stem was six, and these vascular bundles branching and positioned close to the larval chambers (**Fig. 1J, K**).

**Figure 1.**
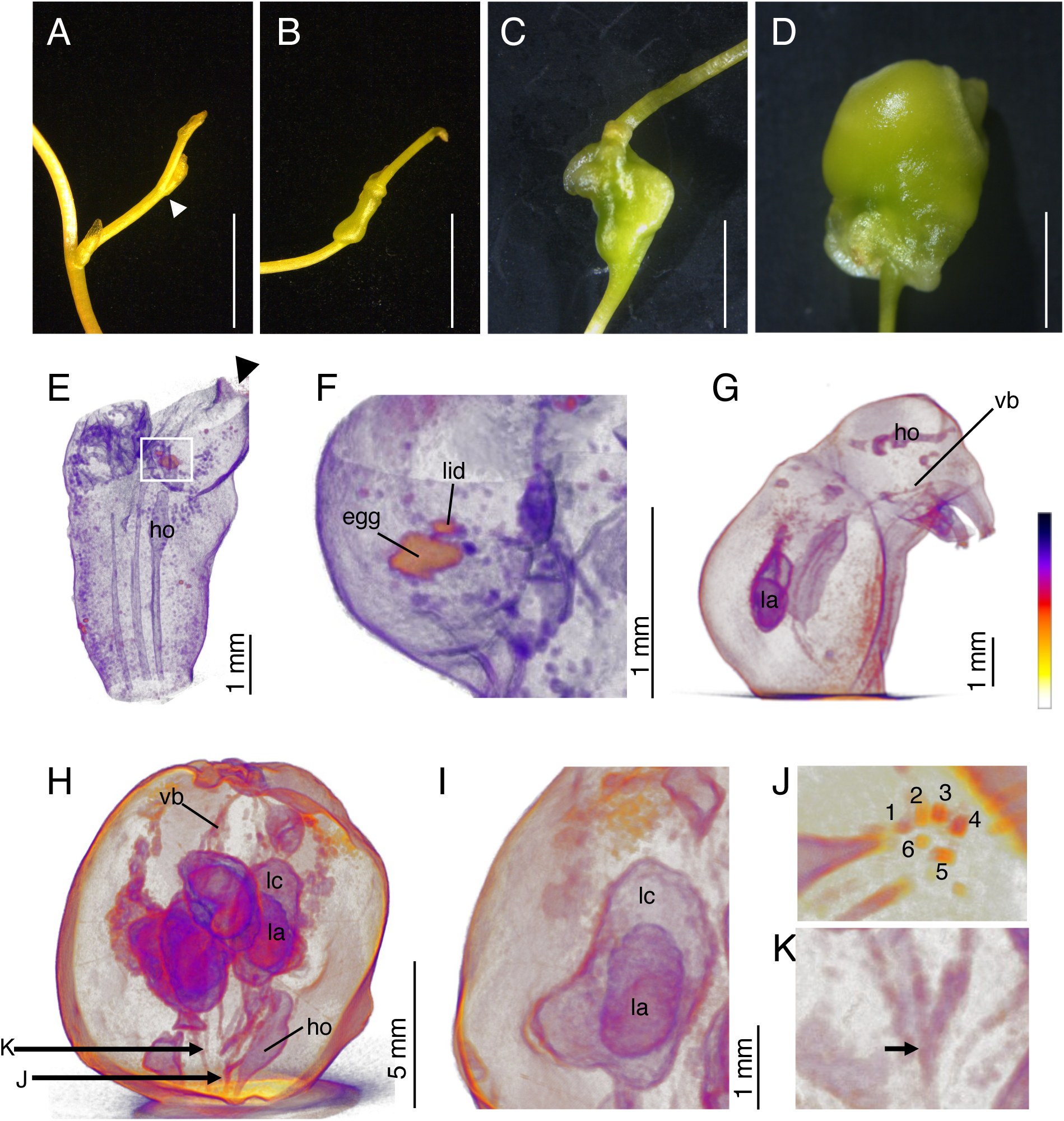
Morphological characteristics of each stage of galls in *C. campestris*. (A-D) Representative image of young node before oviposition (A), Sgall (B), Mgall (C) and Lgall (D). White triangle in (A) indicate node position of *C. campestris* stem. Scale bars indicate 5 mm. (E-I) Representative 3D renderings from X-ray micro-CT scans of Sgall (E), magnified part of Sgall (F), Mgall (G), Lgall (H) and magnified part of Lgall (I). A ‘heat map’ color scale represents X-ray absorption. Same color represents similar x-ray absorption, indicating similar tissue characteristics. Black triangle in (E) indicate the location stem connected. White square indicates magnified region shown in (F). (J-K) Vascular bundles detected in Lgall. Transverse view of the bottom center (J) and branched vascular bundle (K). Arrow indicates branching point. Ho, hole; la, larva; lc, larval chamber; vb, vascular bundle.

### Cellular differentiation depends on the gall growth stages

To examine cellular differentiation in *C. campestris* galls at different stages, we analyzed transverse histological sections (**Fig. 2A-C, Supplementary Fig. S2A-D**). An oviposition hole was observed in the upper part of Sgall as indicated in **Fig. 1E** (white square) (**Supplementary Fig. S2B**). Schizogenous ducts were conspicuous around vascular bundles and newly formed vascular bundles observed around larval chamber (**Supplementary Fig. S2E, F**). Compared to the node, the variation of cell number was high at the Sgall but low at the Mgall stage, indicating rapid cell division occurred at the Sgall stage that slowed at the Mgall stage (**Fig. 2D**). The distribution of cells in transverse sections showed smaller cells densely packed in the central region, with larger cells towards the outer gall region, a pattern consistent across all gall stages (**Supplementary Fig. S2B-D**). However, the size of the parenchyma cells on the outer region increases as the stages progressed (**Fig. 2A-C**). The median cell size per transverse section of Sgall and Mgall were 100 µm^2^ and 240 µm^2^, respectively, indicating an increase in cell size from Sgall to Mgall (**Fig. 2E**).

**Figure 2.**
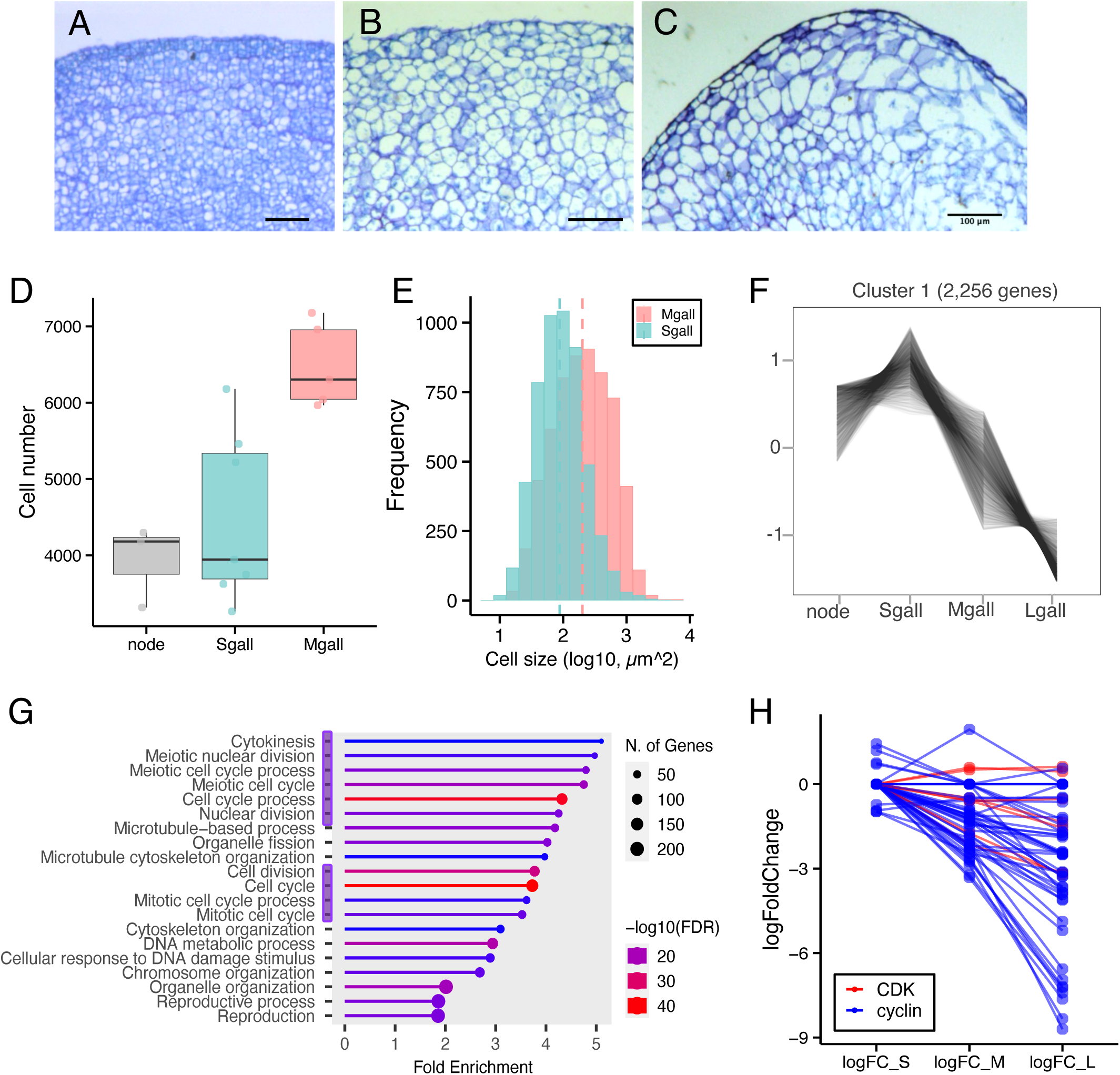
Cell proliferation depends on gall development. (A-C) Representative images of outer layer of transverse paraffine section of Sgall (A), Mgall (B) and Lgall (C). Scale bars indicate 100 µm. (D) Cell numbers in node, Sgall and Mgall. (E) Histogram of the cell size of Sgall and Mgall. Dashed lines indicate median value. *n* = 5,221 (Sgall), *n* = 5,967 (Mgall). (F) Clustering analysis with normalized CPM data by *clust*. Other clusters were shown in **Supplementary Fig. S2**. (G) GO terms of biological process enriched in cluster 1. Color indicates -log10FDR and circle size indicates number of genes. The categories marked in purple are related to cell division or the cell cycle. (H) Gene expression pattern of cyclin and cyclin dependent kinase (CDK). logFC is calculated compared with node.

To understand the molecular mechanisms of *C. campestris* gall development, we conducted transcriptome analysis using three gall stages and unoviposited nodal regions as control. PCA analysis revealed distinct transcriptome profiles for each gall stage and unoviposited nodal regions (**Supplementary Fig. S3A**). Differentially expressed genes (DEGs) were categorized by clustering based on their expression patterns across the developmental stages (**Fig. 2F, Supplementary Fig. S3B**). Gene ontology (GO) enrichment analysis of DEGs in each cluster was performed. In Cluster 1, which showed increased expression at the Sgall stage and decreased expression as the gall developed (**Fig. 2F**), many GO terms related to cell division and the cell cycle were enriched (**Fig. 2G**). Examining the expression patterns of cyclins and cyclin-dependent kinases (CDKs) revealed consistent decreases in expression from the Sgall to Lgall stages (**Fig. 2H**). This is consistent with histological findings that cell division is highly active during the early gall stages, leading to an increase in the number of smaller cells. As the gall matures, cell division decreases, giving way to cell enlargement.

### Expression of photosynthesis-related genes in *C. campestris* galls

Genes showing expression changes were categorized into four clusters besides Cluster 1 (**Supplementary Fig. S3B**). GO enrichment analysis was performed for each cluster and GO terms were enriched in three clusters. In Cluster 2, which consistently showed decreased expression from node to Lgall, GO terms such as DNA replication and ribosome were enriched (**Supplementary Fig. S3C**). Clusters 3 and 4, which showed consistent expression increases from node to Lgall, or sequential increases from node to Mgall with no significant difference between Mgall and Lgall periods, significantly enriched GO terms related to photosynthesis (**Supplementary Fig. S3D, E**). Other GO terms enriched in Cluster 3 and Cluster 4 included defense response genes, namely, autophagy, MAPK signaling pathway, plant-pathogen interaction, and biosynthesis of secondary metabolites which are involved in defense responses (**Supplementary Fig. S3D, E**).

### Expression of flowering related genes in *C. campestris* galls

During the initial stages of gall formation, a shift from normal stem development to gall development was anticipated, resulting in a change in cell fate. We identified 624 upregulated DEGs between Sgall and control (false discovery rate [FDR] < 0.01 and expression-level log2FC > 2). GO enrichment analysis of upregulated genes in Sgall revealed significant enrichment of genes involved in floral organ development (**Fig. 3A**). Additionally, terms such as response to abscisic acid (ABA) and hormone-mediated signaling pathways were also enriched. To understand the expression patterns of flowering-related genes, we particularly focused on the expression changes of class ABCE genes and their upstream regulatory genes, represented in a heatmap. A heatmap of the DEGs involved in floral organ development showed that the expression of many genes increased during the Sgall stage, followed by a decrease (**Fig. 3B**). Especially *AGAMOUS* (*AG*) and *SEPALLATA* (*SEP*) genes were highly expressed (log2FC > 3) at the Sgall stage but decreased in later stages. On the other hand, the expression of *Flowering locus T* (*FT*), a positive regulator of flowering, remained unchanged from node to Sgall but increased during the Mgall stage (**Fig. 3C, Supplementary Fig. S4**). Interestingly, the expression of *Terminal Flower 1* (*TFL1*), a negative regulator of flowering, showed increased in later stages.

**Figure 3.**
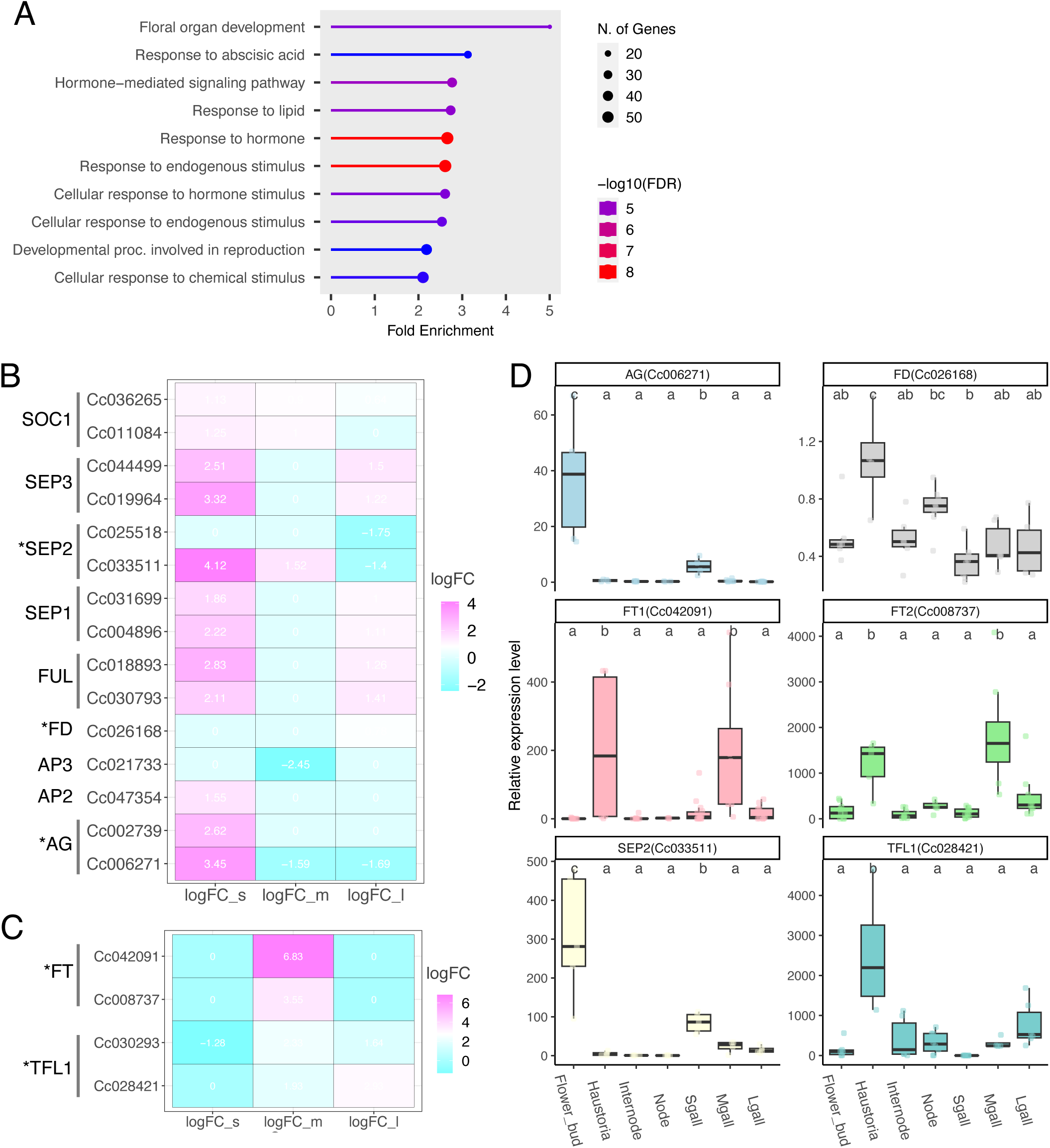
Time-course transcriptome analysis using several stage of *C. campestris* galls. (A) GO terms terms of biological process enriched in upregulated genes of Sgall compared with node (logFC > 2). Color indicates -log10FDR and circle size indicates number of genes. (B-C) Heatmap showing the expression changes of *C. campestris* orthologs of flowering related genes depends on gall development based on the RNA-seq data. The heatmaps present the logFC values, relative to the previous stage of tissue (logFC_s means Sgall vs node, logFC_m means Mgall vs Sgall, logFC_l means Lgall vs Mgall. Asterisks indicate genes that examined the relative expression levels by qRT-PCR in Fig. 3D. (D) Relative expression levels of flowering related genes measured by qRT- PCR in several tissues of *C. campestris*. Expression levels with three to seven biological replicates were normalized with respect to that of *CcACT8*. Statistically significant differences are indicated by different letters above the boxes (ANOVA with Tukey’s HSD test, P < 0.05).

The expression levels of these key flowering-related genes were verified by quantitative reverse transcription PCR (qRT-PCR). In qRT-PCR, several tissues such as internode, haustoria, and flower buds were used as samples in addition to the galls. The results showed that *AG* and *SEP2* were highly expressed in flower buds (**Fig. 3D**) where both genes functioned. An increase in the expression of these genes during the Sgall stage was observed compared to the node or other gall stages. Consistent with previous findings (Mäckelmann et al., 2024), *FT1* and *FT2* were highly expressed in the haustoria. Additionally, increased expression of these genes was observed during Mgall stage (**Fig. 3D**). Interestingly, *TFL1* was highly expressed in the haustoria, co-localizing with high *FT1* and *FT2* expression. The expression of *FD*, a gene known to interact with both *FT* and *TFL1*, was elevated in haustoria but was also ubiquitously across other tissues as previously reported (Mäckelmann et al., 2024). These results indicated that downstream genes in the flowering pathway, like *AG*, were predominantly expressed at the initial gall developmental stage. In contrast, upstream genes of the flowering pathway, like *FT*, were more actively expressed during the later stage of gall maturation. The differential expression patterns of these genes appear to correlate with the different stimuli arising from the adult and larval stages of *S. madaranus* (**Supplementary Fig. S5**).

### Spatial expression patterns of flowering-related and stem cell niche marker genes in *C. campestris* galls

To understand the spatial expression patterns of genes involved in the flowering pathway and their contribution to gall development, we conducted *in situ* hybridization analysis using Sgall and Mgall. *AG* was predominantly expressed in parenchyma cells at Sgall stage, but this expression pattern was restricted to the areas surrounding vascular tissues at the Mgall stage (**Fig. 4A-D**). Conversely, *FT2* was faintly expressed at the Sgall stage but showed strong expression in the central areas around the larval chambers in Mgall tissues (**Fig. 4E-H**). Gall development, a form of *de novo* organogenesis, is thought to require stem cells as a foundational element (Schultz et al., 2019; Hirano et al., 2020). We found that the stem cell niche marker gene, *PLETHORA* (*PLT*) was expressed specifically at Sgall according to the transcriptome analysis (logFC = 6.61 at Sgall compared to node) and it was validated by qRT-PCR (**Supplementary Fig. S6A, B**). *PLT* signals were observed throughout the Sgall tissue (**Fig. 4I, J**), indicating active cell division. A strong signal of *PLT* was also observed in the central area and surrounding vascular tissues in Mgall (**Fig. 4K, L**). These results indicated that both the majority of the Sgall tissue and the central area of Mgall exhibited stem cell-like characteristics.

**Figure 4.**
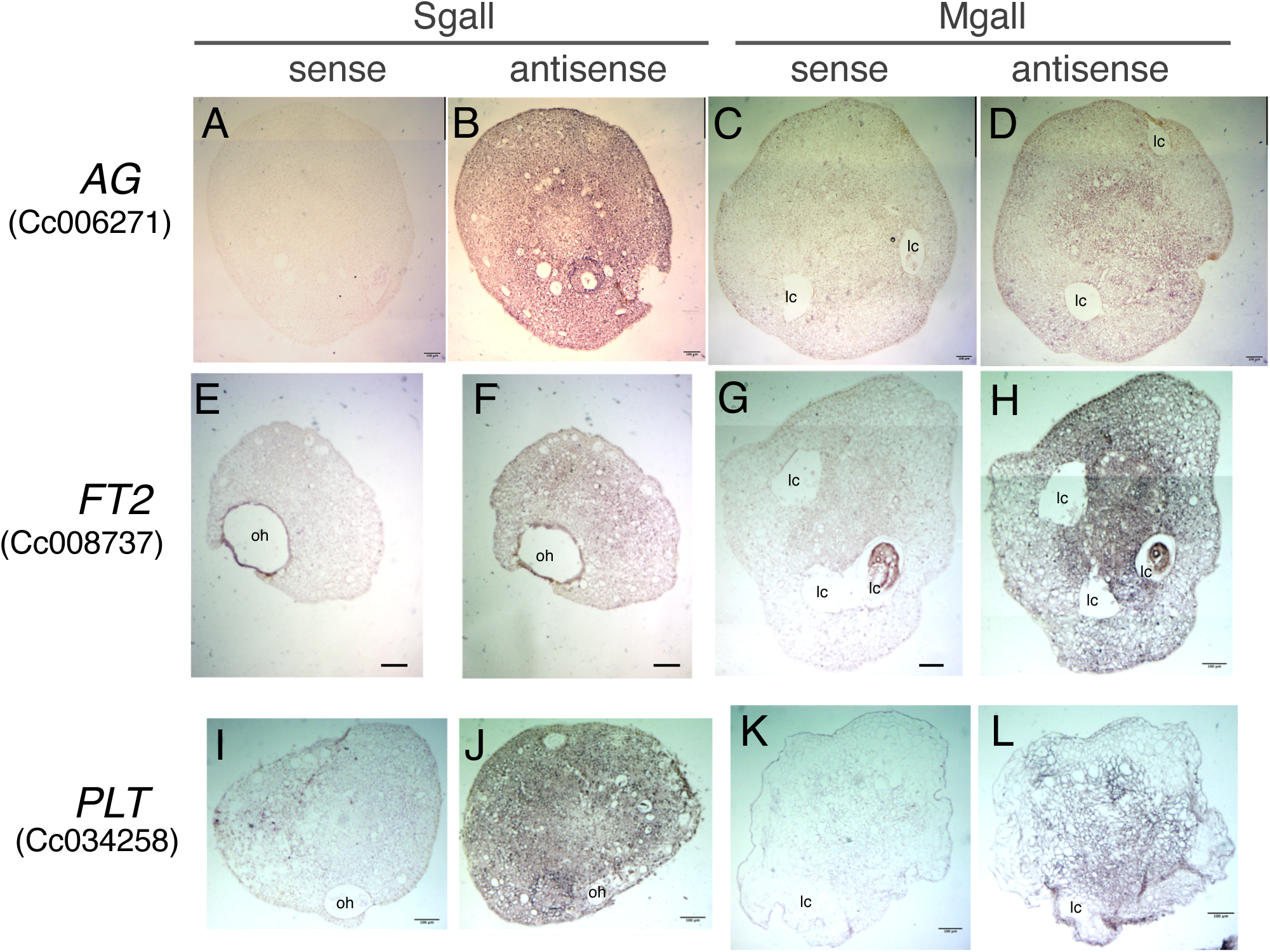
Expression patterns of flowering-related and stem cell niche marker genes in galls. (A-I) *in situ* hybridization using *CcAG* (A-D), *CcFT2* (E-H), *CcPLT* (I-L) probes on Sgall and Mgall samples. Scale bars indicate 100 µm. oh, oviposition hole; lc, larval chamber. *AG*, *AGAMOUS*; *FT2*, *Flowering locus T2*; *PLT*, *PLETHORA*.

### Upregulation of homeobox genes in early gall stage

To identify the meristematic marker genes comprehensively other than *PLT*, GO enrichment analysis was performed. The GO term of meristem development (GO:0048507) was enriched in the upregulated DEGs in Sgall involving 23 genes (**Supplementary Table 7**), including homeobox genes, *WUSCHEL-related homeobox 2* (*WOX2*) and *ATHB-14*. As homeobox genes like *SHOOT MERISTEMLESS* (*STM*) and *WUSCHEL* (*WUS*) are known to induce ectopic meristems (Gallois et al., 2002), we analyzed the expression pattern of identified homeobox genes across the developmental stages. The results indicated that *WOX2* and 24 other homeobox genes were significantly upregulated in the Sgall stage (logFC > 1.0), with the highest being *Cc006977* at logFC is 5.16 (**Supplementary Table 9**). Conversely, homeobox genes upregulated in the Mgall and Lgall stages were predominantly from different classes of those of Sgall, and their numbers decreased from 25 in the Sgall stage to 8 in Mgall and 5 in Lgall.

### Impact of shading treatment on gall development

To assess the impact of photosynthesis on gall development, we conducted a shading experiment (**Fig. 5**). In this experiment, we measured the size of galls, the amount of chlorophyll and starch, and the developmental stages of the insects inside galls under both shaded and unshaded conditions. To ensure equal nutrient and water absorption, we adjusted the number of *C. campestris* stem and the number of haustoria on the host to be the same. The shaded galls show a significant difference in color and size compared to the control (**Fig. 5A, B**). Regardless of whether there was one or two larvae inside of the galls, the gall volume under shaded conditions was consistently smaller compared to the control (**Supplementary Fig. S7A**). Additionally, there were significant differences in the amounts of chlorophyll *a* and *b* but not in carotenoids (**Fig. 5C**). This result supported the color change after shading treatment. Unexpectedly, the starch amounts in the galls between the shaded group and the control were not significantly different (**Supplementary Fig. S7B, C**). To evaluate the effect of shading treatment on insect development, we checked the stage of *S. madaranus* inside the galls. In shaded galls, the proportion of larvae was significantly higher (71.4%) while the proportion of pupae was lower (28.6%) compared to the control (χ^2^ =5.73, P<0.05, **Fig. 5D**). The results showed that shading reduced chlorophyll levels and affect to gall size and larval growth.

**Figure 5.**
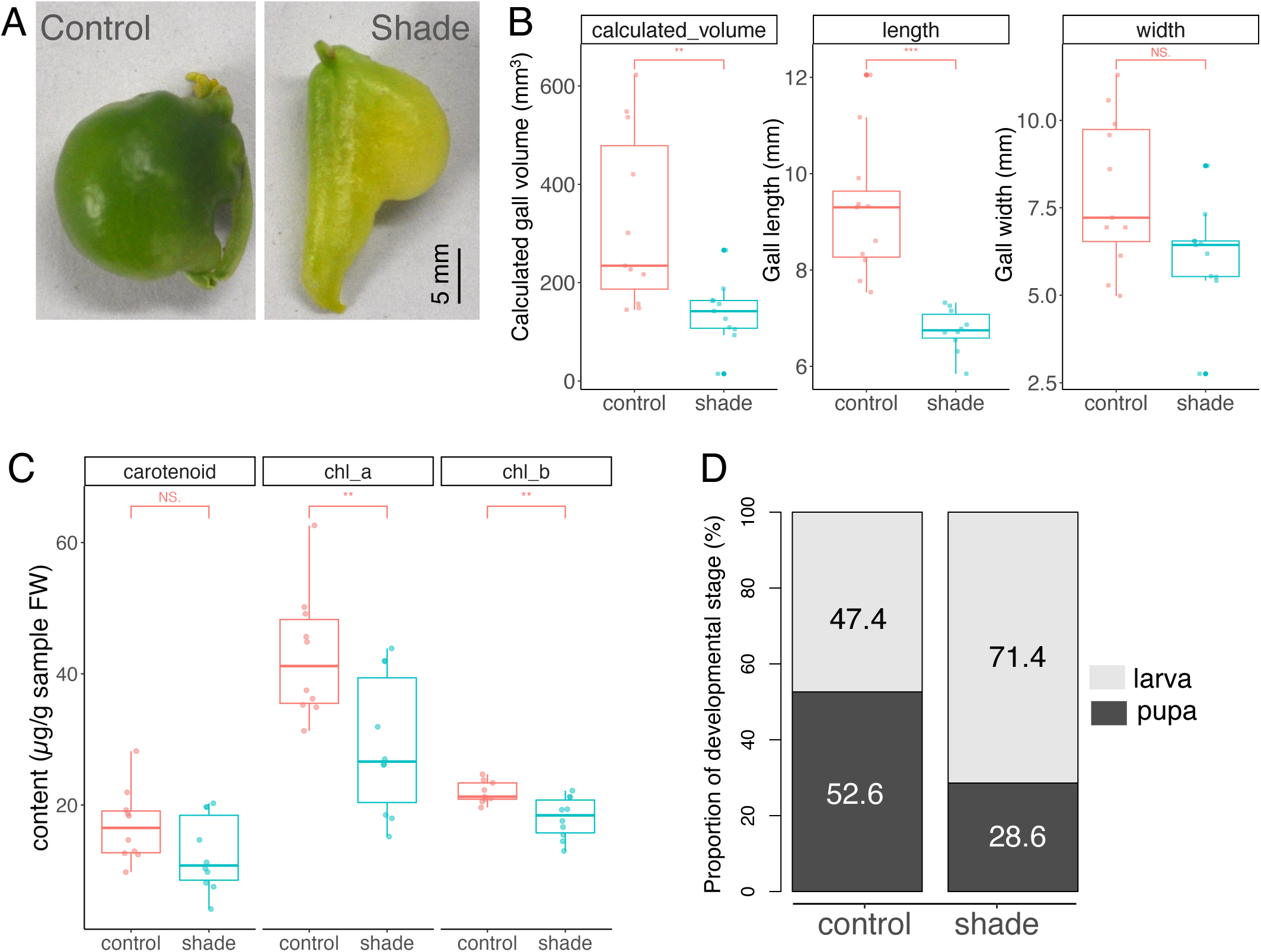
Effects of shading in developing galls. (A) Galls with or without shading treatment. (B-C) Effect on gall development by shading. Calculated gall volume (B), chlorophyll contents (C). **P < 0.01, ***P < 0.001, NS, not significant difference; Wilcoxon rank sum test with Bonferroni correction. (D) Developmental stage of *S. madaranus* in the galls of both treatments. Significance of the proportion difference between the treatments was assessed using a chi-square test.

## Discussion

Gall development involves the complex interplay between insect-derived molecules and plant developmental pathways. Previous studies have highlighted the activation of flowering-related genes during gall formation. For instance, in *Rhus javanica* galls induced by the aphid *Schlechtendalia chinensis*, genes like *APETALA1* (*AP1*, class A), *SEP* (class E), and *LEAFY* (*LFY*) are upregulated (Hirano et al., 2020). Similarly, in *Artemisia montana* galls, genes such as *AP1* (class A), *PI* (class B), and *SEP* (class E) are expressed in initial stage of galls (Takeda et al., 2019). Schultz *et al*. (2019) conducted RNA-seq analysis across four developmental stages of *Phylloxera*-induced galls on grapevine leaves (Schultz et al., 2019). They found that *TFL1* expression increased in early galls, while *AG* and *SEP* (class C and E) showed higher expression in later stages. In this study, we showed that the genes in the flowering pathways were also activated during stem-derived gall formation in *C. campestris.* However, while there were some similarities, the activated genes were somewhat different from those identified in the previous studies. Our time-course RNA-seq revealed increased *AG* and *SEP* expression in the Sgall stage, similar to those in *R. javanica* and *A. montana* leaf-derived galls, but no increase in *LFY* expression. The combination of *AG* and *SEPs* is required to produce a carpel (Ferrándiz et al., 2010), suggesting that the transition from normal stem to carpel-like characteristics might cause gall initiation in *C. campestris* gall. Unlike grapevine leaf galls, *TFL1* expression increased in later gall stages, alongside *FT* expression in *C. campestris* galls. Genes controlling *FT* expression in *A. thaliana*, such as *CONSTANS* (*CO*) and *GIGANTEA* (*GI*) showed no expression changes, similar to other plant galls. This difference could be attributed not only to the location of gall formation (leaf vs. stem), but also to variations in the insect’s reproductive modes (asexual reproduction inside of the gall vs. sexual reproduction), feeding style on the plants (sap-sucking vs. biting), and gall induction manner (feeding vs.feeding and oviposition).

Our study also supports the idea that gall induction and growth are influenced by specific insect developmental stages; Oviposition by adult females triggers early gall development, while larvae feeding drives later-stage gall enlargement (Ushima et al., 2024). In this study, we demonstrate that the expression of downstream flowering pathway genes increases during the early gall stage, which is responding to the stimuli by adult females (**Supplementary Fig. S5**). We also show that induction of upstream flowering genes occurs during the later stage of gall formation induced by larvae. This suggests that the insect might activate the flowering pathway at unconventional stages, potentially rerouting the typical flowering cascade through effector molecules to influence gall development. However, it remains unclear whether this phenomenon is a direct or indirect response to *S. madaranus*. Several reports indicate that ABA accelerates flowering by promoting the expression of flowering-related genes (Domagalska et al., 2010; Riboni et al., 2013; Riboni et al., 2014). The elevation of ABA response genes in Sgall might explain the upregulation of flowering-related genes. Moreover, biotic stress including herbivory often triggers changes in gene expression, including flowering genes (Takeno, 2016), which could relate to *FT* expression in Mgall and Lgall stages. To further explore this, investigating the role of ABA and other stress hormones could provide valuable insights into how insect-induced gall formation affects flowering pathways. Notably, RNA interference (RNAi) has been effective for gene function analysis in *S. madaranus* (Ushima et al., 2024), paving the way for more detailed molecular analysis of the insect’s role in gall formation.

Our findings on the cellular dynamics of gall development suggest parallels to fruit growth. In many fruits, rapid cell division in the early stages is followed by cell enlargement (Gillaspy et al., 1993; Bertin et al., 2003). A similar pattern is observed in *C. campestris* galls. In the Sgall stage, rapid cell division occurs, accompanied by increased expression of *PLT* and many meristem-related homeobox genes. Earlier studies using leaf-derived galls have similarly shown that key regulators of stem cell development, such as homeobox genes, play crucial roles in gall formation (Schultz et al., 2019; Hirano et al., 2020, Takeda et al., 2021). In this study, the upregulation of *PLT* and meristem-related homeobox genes during the Sgall stage indicates that gall formation involves both the induction and maintenance of stem cells, highlighting a common mechanism shared between leaf-derived and stem-derived galls. However, as the gall develops to the Mgall and Lgall stages, cell enlargement might become more prominent, suggesting a shift from active cell division to cell expansion, much like the developmental trajectory seen in fruit growth. The formation of meristematic tissue around the larval chamber acts as a continuous source of cells for nutritive tissue and adjacent regions. We observed some homeobox genes highly expressed in the late stage of *C. campestris* galls and verified that *PLT* kept expressing in the central area of Mgall. Schultz *et al*. (2019) also reported increased expression of meristem maintenance genes in later stages (Schultz et al., 2019), which suggests that different meristem-related genes may be upregulated depending on the developmental stage of the insect inside, thereby driving plant cell proliferation.

Previous studies showed a decrease in photosynthesis-related gene expression in leaf-derived galls compared to leaves (Nabity et al., 2013; Takeda et al., 2019; Hirano et al., 2020) and even in stem-derived galls (Takeda et al., 2024). In contrast, our RNA-seq analysis on stem-derived galls of parasitic plant *C. campestris*, which typically show little photosynthetic activity (Pattee et al., 1965; Sherman et al., 1999), revealed an increase in photosynthesis-related gene expression as the galls developed. This was associated with the findings that *Cuscuta* galls induced by *Smicronyx* exhibit higher chlorophyll and chloroplast concentrations, along with increased photosynthetic activity compared to stems (Anikin et al., 2017; Zagorchev et al., 2018; Murakami et al., 2021; Zagorchev et al., 2021). But why does chlorophyll accumulation occur in *C. campestris* galls? McNeal et al. (2007) reported chlorophyll accumulation during ovule development in *Cuscuta* genus, contributing to the synthesis of lipids serving as energy reserves for seedlings. Given the low availability of lipids from the host’s vascular extracts and the high demand for lipid production during fruiting, this efficient lipid synthesis pathway offers a more plausible explanation for the preservation of a photosynthetic apparatus in *Cuscuta* (McNeal et al., 2007). Additionally, the enrichment of the GO term "response to lipid (GO:0033993)" at all stages of gall development suggests that lipids are accumulating in the galls of *C. campestris* (**Supplementary Table 7**). Under shaded condition, chlorophyll content decreased in *C. campestris* gall as well as in other plants (Pružinská et al., 2005; Song et al., 2014), and the development of *S. madaranus* within the galls slowed. However, there was little difference in starch content between the control and shaded galls. This suggests that the primary role of photosynthesis in the galls is not only to supply starch to the larvae but potentially to produce other products. Another study has also shown that the growth of the larvae inside the shaded galls was substantially reduced (Haiden et al., 2012). The authors suggested that photosynthesis at the gall may contribute to oxygen provision in addition to photosynthates. In the case of *C. campestris* gall, chlorophyll may contribute to supply lipids or potentially oxygen to the larvae within the closed galls for promoting normal development. These aspects will be further explored in future research. Additionally, research on tomatoes, which belong to the same order *Solanales* as *C. campestris*, has shown that photosynthesis-related genes are upregulated in the locule, which is located at the center of the tomato fruit (Lemaire-Chamley et al., 2005; Lytovchenko et al., 2011). Chlorophyll accumulation and photosynthesis in tomato fruits play an important role in fruit metabolism and development (Dong et al., 2024). These findings suggest that the signaling pathways involved in flowering, fruit development, and photosynthesis observed in *C. campestris* galls may reflect certain traits that are present in *Solanales* fruits.

This study clarifies that the developmental pathway of *C. campestris* galls at the gene expression, cellular, and physiological levels partially follows the fruit developmental pathway. This provides a fascinating example of how gall-inducers can co-opt host resources and genetic pathways to support their growth and reproductive success. These findings underscore the complexity of plant-insect interactions and offer new insights into the adaptive strategies of gall-inducing insects.

## Materials and Methods

### Plant and insect materials and growth conditions

A laboratory strain of *S. madaranus* was collected from Neagari Nomi City, Ishikawa, Japan. It was maintained on *C. campestris* parasitizing *Nicotiana benthamiana* in the cargo box and placed at growth room at 25 °C in a long day condition (14h Light/10h Dark) as described (Murakami et al. 2021). In this paper, we refer to the “initial gall” (Murakami et al. 2021) as Sgall, which appears approximately two days after the insect introduction. Mgall and Lgall were defined as stages 6 and 10 days after Sgall formation (**Supplementary Fig. S1A**). In Sgall, eggs are always present, but no larvae. In Mgall and Lgall, larvae are present (Ushima et al., 2024). The pictures of each stage of gall and *C. campestris* node were taken by Leica MZ10F (Wetzlar, Germany).

### micro-CT imaging

Several stages of *C. campestris* galls were collected from the lab conditions on December 23, 2022, for micro-CT scanning. Galls were fixed in 4% paraformaldehyde (PFA) solution in 1X phosphate-buffered saline （PBS), and after subsequent immersion in 70% (v/v) and 35% (v/v) ethanol, distilled water, and 25% (v/v) Lugol’s solution as described previously (Maeno and Tsuda, 2018). The samples were scanned using ScanXmate L090T (Comscan techno, Japan) with a tube voltage of 25 kV and tube current of 170 µA. Visualization and measurement of the micro-CT images were conducted using Volume Viewer of *Fiji* ver. 2.15.1 (Schindelin et al., 2012). *Fiji* can apply color mapping to the dataset to highlight the different regions as shown in **Fig. 1E-K**.

### Histological analysis using paraffin sections

About three-weeks-old tobacco (*N. benthamiana*) plants were attached with 5 cm long *C. campestris* vine. Adult *S. madaranus* weevils were infested to *C. campestris* for inducing galls. Two days after inoculation of the weevils, Sgalls were collected and fixed in the fixative solution consisting of 4% PFA in PBS under vacuum conditions until the samples sank to the bottom of the tube. After fixation, the samples were dehydrated through a graded ethanol series, (50, 70, 80, 90, 95 and 100% ethanol) for 45 minutes each and then followed by Lemosol (Wako, Japan) in ethanol series (2:1, 1:1, 1:2 and 100% lemosol) for 45 minutes each. The tissues were then infiltrated with Paraplast Plus (Sigma-Aldrich, USA) at 60°c with overnight incubation. After the tissue was embedded in paraffin, it was mounted on a wooden block, which was then clamped onto a microtome for section cutting. The embedded samples were cut to 10 µm thickness to form ribbons by MICROM HM325 rotary microtome, flattened fully on adhesive glass slides (Matsunami, Japan) placed on a hot plate at 42℃, and then dried. The sections were deparaffinized in 100% Lemosol solution and rehydrated through a graded ethanol series. The sections were stained with 1% Safranine and 0.1% Toluidine blue O for histological observation.

### Histological analysis using Technovit resin

Tissues were fixed in 500 µL of 4% PFA solution for two hours. Following fixation, the samples were dehydrated sequentially in graded ethanol solutions (70%, 80%, 90%, 95%, and 100%). The dehydrated samples were then placed in a tube containing 500 µL of 100% ethanol and agitated on a rotator at approximately six rotations per minute, with 100 µL of Technovit 1 solution (Technovit 7100 + hardener I) (Nisshin EM Co., Ltd., Japan) added every 10 minutes until a total volume of 1000 µL was reached. The Technovit/ethanol mixture was then removed, and 500 µL of fresh Technovit 1 solution was added to the sample, followed by a 1-hour incubation. Post-incubation, the samples were transferred to embedding dishes, and 1 mL of Technovit II solution (Technovit 7100 + hardener II) (Nisshin EM Co., Ltd., Japan) was added. The samples were left overnight at 4°C, covered with parafilm, to allow the resin to harden. Once hardened, the resin blocks were removed from the embedding dishes, trimmed, and attached to support blocks using instant adhesive. The samples were then clamped onto a microtome for sectioning. The embedded samples were sectioned at a thickness of 10 µm using a MICROM HM325 rotary microtome. The sections were then placed on adhesive glass slides (Matsunami, Japan) and dried on a hot plate at 42°C. The sections were subsequently stained with 1% safranin for 15 seconds, washed under running water, dried, and observed under a microscope for further analysis.

### Cell segmentation

Cell size of transverse gall section was quantified using *Fiji* ver. 2.15.1 (Schindelin et al., 2012) using safranin-stained sections. Cell identification using *Find Maxima* function under the *Process* menu. Settings included checking the *Preview* point selection option, setting the *Output* type to *Segmented particles*, and enabling the exclusion of cells touching the edge. An automated segmented cell image was then generated and saved for analysis. For analysis, measurement settings were configured by navigating to *Analyze* and selecting *Set Measurements*. Necessary measurement options such as area, shape descriptors, display label, mean gray value, and perimeter were chosen. For particle analysis, the *Analyze Particles* function under *Analyze* was used. Settings were adjusted to display outlines and results. This process resulted in an image with detailed information on each cells size and shape metrics, ready for analysis.

### Transcriptome analysis

Total RNA was extracted from different developmental gall stages (unoviposited nodal region including 1 cm of stems at the node, Sgall, Mgall and Lgall as described above) by Maxwell RSC Plant RNA kit (Promega, USA) according to the manufacturer’s instructions. Total RNA integrity was quantified using an Agilent 2100 Bioanalyzer (Agilent Technologies, USA). RNA-seq libraries for Illumina sequencing were constructed with TruSeq kit (Illumina, USA) and the qualities of libraries were checked on an Agilent 2100 Bioanalyzer. Libraries with three biological replicates per stage were sequenced on a HiSeq2000 Illumina Sequencer at Macrogen co. (Tokyo, Japan). A total of 621,387,924 raw sequencing reads were generated (10–17 million read pairs per sample; **Supplementary Table 1**) and used for gene expression profiling. Raw reads were mapped to the published genome of *C. campestris* (Vogel et al., 2018) with Hisat2, and subsequently read count table was created by using StringTie and prepDE.py (Pertea et al., 2016).

### Functional annotation and GO enrichment analysis of RNA-seq data

For *C. campestris* genes, since many genes are not functionally annotated in the current published *C. campestris* genome (Vogel et al., 2018), we did blastp to the *Arabidopsis thaliana* genes for annotated *C. campestris* transcriptome with e-value < 1e-04. We also used eggNOG-mapper (Cantalapiedra et al., 2021) for identifying hypothetical proteins. TAIR ID hits were used for GO Enrichment Analysis on http://geneontology.org/ for gene clusters and modules. ShinyGO ver. 0.80 was used for visualization of GO analysis results (Ge et al., 2019). Detection of the differentially expressed genes were performed by edgeR (Robinson et al., 2010) with FDR < 0.01 and log2FC > |2|. Clustering analysis was done by using *clust* (Abu-Jamous and Kelly, 2018) with default option and clustered genes based on read count. The genes including in each cluster are listed in **Supplementary Table 2-6**.

### Expression analysis by qRT-PCR

Several stage of gall samples as indicated Sgall, Mgall and Lgall were collected from lab-grown samples. Node including stem axis was collected one week after parasitism induction on *N. benthamiana*. Flower bud, internode, and haustoria samples were collected one month after parasitism induction. Tissues were frozen in liquid nitrogen immediately after collection and homogenized by Tissue Lyser II (QIAGEN, Netherlands). Total RNA was isolated using a Maxwell RSC Plant RNA Kit (Promega, Madison, WI). RNA concentration and purity were assessed using a Nano-Drop® ND- 2000 spectrophotometer (Thermo Scientific, USA). For synthesizing first strand cDNA, we used 1 µg of total RNA as a template using the oligo-dT primer supplied with SuperScript^TM^ IV kit (Thermo Scientific, USA). Quantitative real time (qRT) PCR of the target genes was performed using SYBR™ Green qPCR Master Mix (Thermo Scientific, USA) and LightCycler^®^ 480 Instrument (Roche, Switzerland) with specific primer sets, as shown in **Supplementary Table 10**. The PCR temperature profile was 95°C for 1 min, followed by 40 cycles of 95°C for 15 s, 60°C for 1 min as 2-step PCR. The dissociation stage was performed at 95°C for 15 s, 60°C for 1 min, followed by a slow ramp to 95°C. *CcACT8* was used as an internal control for normalization. Quantitative PCR and dissociation curve analyses were performed on three to seven biological replicates per tissue using a standard curve method.

### *in situ* hybridization

RNA probes for *in situ* hybridization were produced using *CcAG*, *CcFT2* and *CcPLT* partial sequences cloned in pENTR-D-Topo vector using primers listed in **Supplementary Table S10** followed as manufacture’s protocol. The labeled RNA probes were synthesized using *in vitro* transcription in the presence of Digoxigenin-label by RNA polymerases T7 or SP6 provided with DIG RNA labeling kit (Roche Diagnostics, Switzerland). The tissue sections of 10 µm thickness were sectioned by MICROM HM325 rotary microtome followed by deparaffinization with 100% lemosol for 10 mins and hydrated in graded ethanol series. Further hybridization procedure was performed according to a previously published protocol (Shimizu et al., 2018). Nitroblue tetrazoliumchloride (NBT) (Roche Diagnostics, Switzerland) and 5-bromo-4-chloro-3- indolyl phosphate (BCIP) (Roche Diagnostics, Switzerland) were used as substrates and incubated for 10 minutes or till positive signals on sense/anti-sense probes are seen. After detecting the signals, Immuno In Situ Mounting Medium (Funakoshi, Tokyo, Japan) was added for mounting. Images were obtained using Leica microscope. The captured images were processed using Leica LAS X.

### Shading experiment

Galls finding after two days of Sgall were used for the shading experiment. The stem of 5 mm above and below the galls were covered with foil as a control, while the entire galls, including the stems above and below the galls, were covered with foil as the shading treatment. After 12 days of foil covering, the galls’ pictures were taken after removing foils, then the length and width of the galls were measured using ImageJ (ver 1.51n). Subsequently, the galls were bisected with a razor blade, revealing either one or two larvae within each gall. The developmental stages of the larvae were then recorded. One-half of the bisected galls were used for chlorophyll quantification, and the other half for starch quantification. The calculation of gall volume and the quantification of chlorophyll and starch were followed the methods in the previous study (Murakami et al., 2021). The estimation of total carotenoid was done as described in previous study (Wellburn et al. 1994).

### Data availability

All datasets supporting the results and conclusions of this manuscript are included in the article and supplementary files. Materials generated in the study are maintained at Nagoya University. All reads generated for RNA-seq (PRJDB18311) have been uploaded to the DNA Databank of Japan (DDBJ) repository.

### Quantification and statistical analysis

Statistical analyses and graphical illustrations were all performed using the statistical software R v.4.3.1 with the interface Rstudio v2023.06.1 and the R packages, rstatix, tidyr, ggpubr, ggplot2, purrr, and multcompView. For differences in relative gene expression level, we performed one-way ANOVA followed by a post-hoc analysis with Tukey’s honest significance difference test. For differences in gall size, carotenoid, chlorophyll, and starch content between control and shading treatment, we performed the Wilcoxon rank sum test with Bonferroni correction. The chi-square test was used to examine the difference in the proportion of larvae and pupae in the galls between control and shading treatment.

## Acknowledgments

We thank Mrs. K. Sano, S. Uda, R. Sugimoto and Y. Sano for keeping materials. We thank Dr. T. Oi for teaching how to use X-ray micro-CT machine, Dr. R. Yokoyama for setting up growing condition, and Dr. T. Makino for his fruitful comments. Transcriptome analyses were partially performed on the NIG supercomputer at the National Institute of Genetics.

## Author Contributions

NJU, TT and KBU designed this study and wrote the manuscript. NJU performed histological analysis including *in situ* hybridization. RU, TT and KBU performed RNA-seq analysis. KBU conducted micro-CT analysis. YY conducted qRT-PCR. All authors have reviewed and commented on the manuscript.

---

**Supplemental Figure S1.**
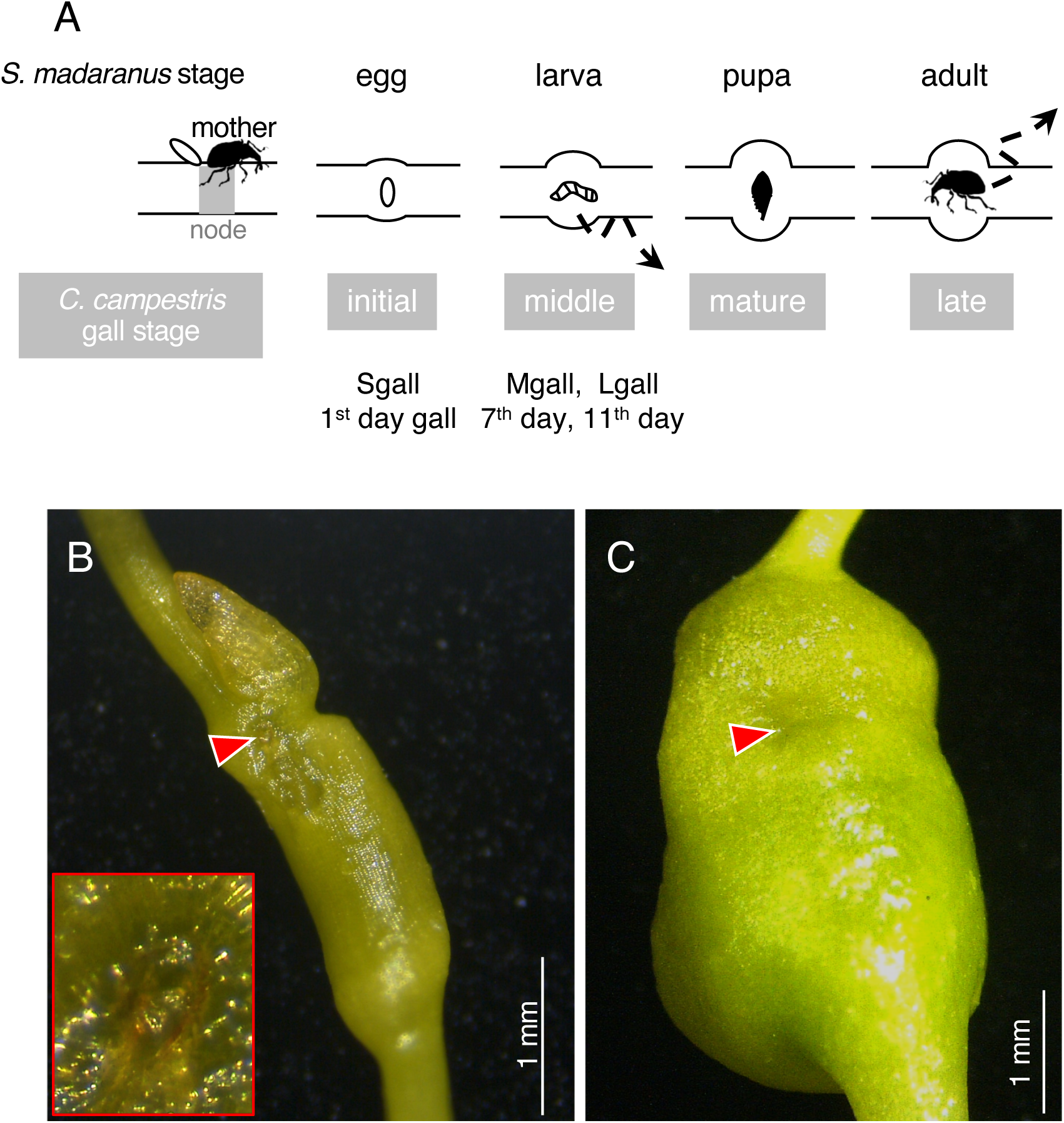
Life cycle of *Smicronyx madaranus*. (A) Mother insects lay eggs at the young node of *C. campestris* stem. Depending on environmental conditions, *S. madaranus* spends its entire lifecycle within the gall from egg to adult, or it emerges from the gall at the final larval stage to pupate in the soil based on our observation. We called Sgall, Mgall, and Lgall that are in the 1st, 7th and 11th day gall stages. respectively. (B) Oviposition hole at Sgall. Red arrowhead and the magnified picture surrounded by red indicate the oviposition hole. (C) Oviposition hole was hardly observed at Mgall stage. Supplementary Figure S2. Complete transverse sections at different stage of C. campestris galls. Node before oviposition (A), Sgall (B), Mgall (C) and Lgall (D). (E-F) Transverse sections of Mgall (E), and Lgall (F). oh, oviposition hole; lc, larval chamber; pa, parenchyma cells; ph, phloem; vb, vascular bundle; xy, xylem. Asterisks indicate schizogenous ducts. Arrows in (F) indicate newly formed vascular bundles. Scale bars are 100 μm.

**Supplementary Figure S2.**
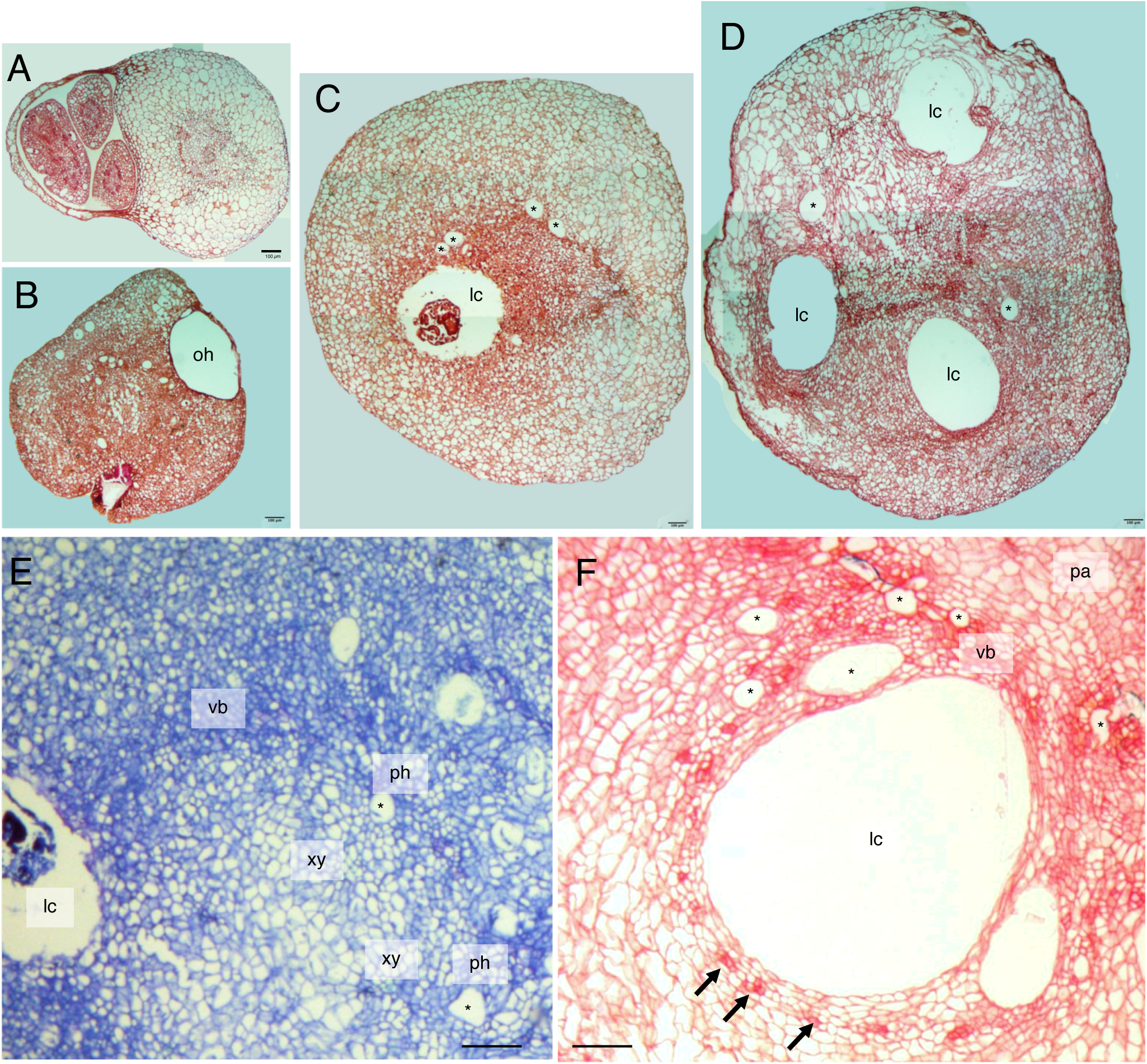
Complete transverse sections at different stage of C. campestris galls. Node before oviposition (A), Sgall (B), Mgall (C) and Lgall (D). (E-F) Transverse sections of Mgall (E), and Lgall (F). oh, oviposition hole; lc, larval chamber; pa, parenchyma cells; ph, phloem; vb, vascular bundle; xy, xylem. Asterisks indicate schizogenous ducts. Arrows in (F) indicate newly formed vascular bundles. Scale bars are 100 μm.

**Supplemental Figure S3.**
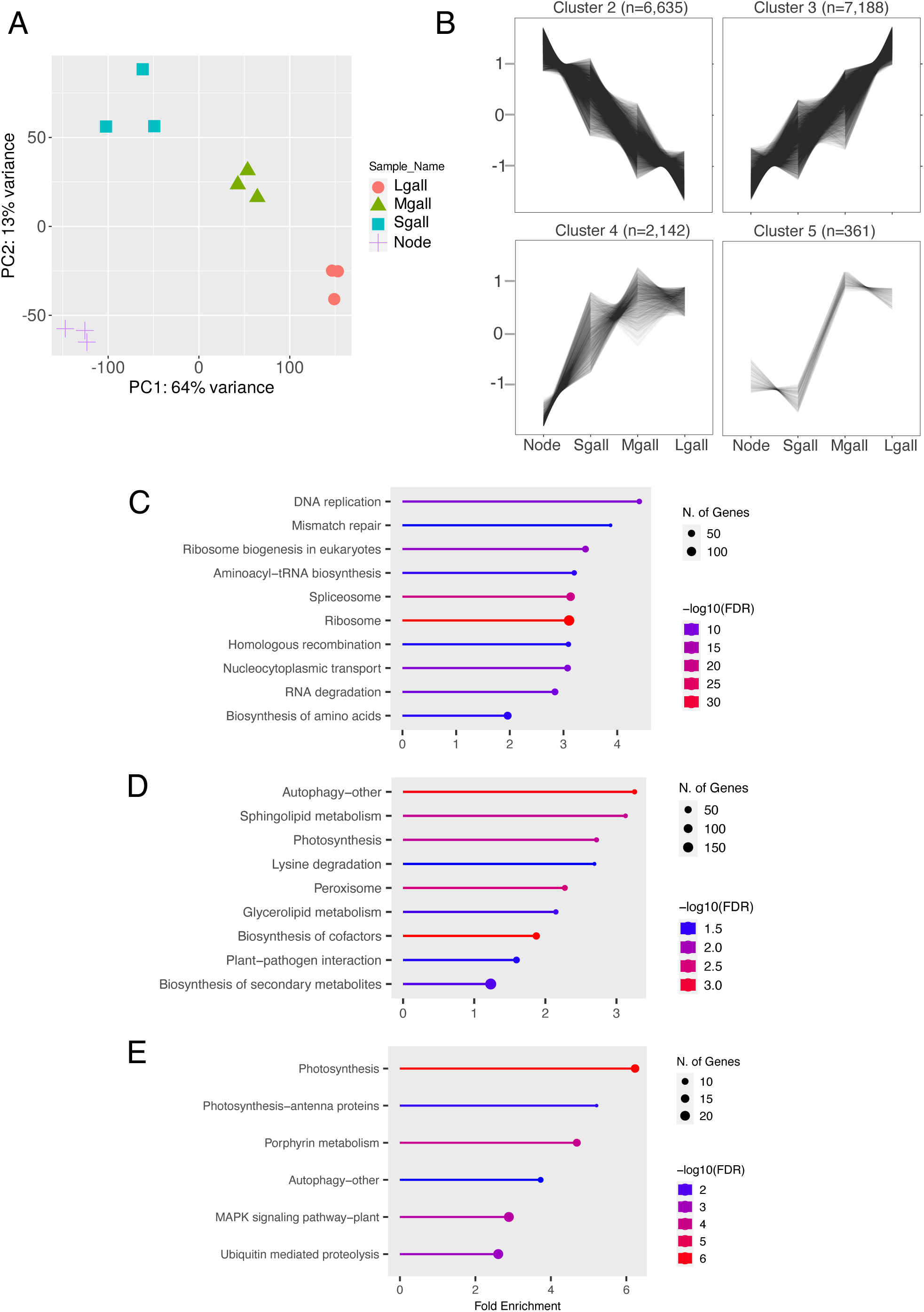
Summary of RNA-seq data using several stage of *C. campestris* galls. (A) PCA analysis on all the samples. (B) Clustering analysis with normalized CPM data by *clust*. (C-E) KEGG pathway enrichment analysis using cluster 2 (C), cluster 3 (D), and cluster 4 (E). No GO term was enriched in cluster 5. FDR < 0.05, Top 10 pathways are represented.

**Supplemental Figure S4.**
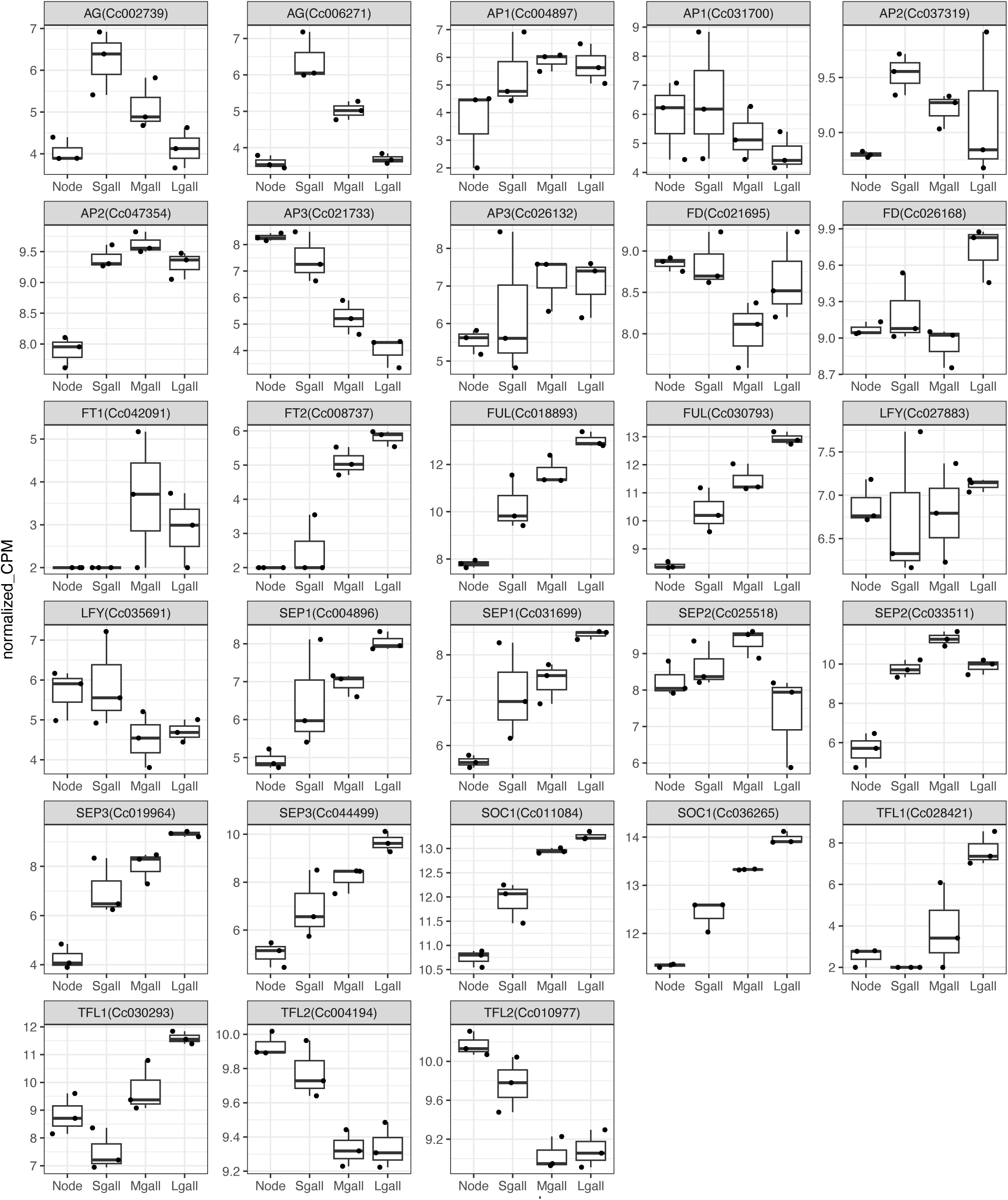
Flowering related genes expression levels in node, Sgall, Mgall, and Lgall quantified as normalized counts per million (CPM). CPMs are calculated by normalizing the read counts by the total counts per sample.

**Supplemental Figure S5.**
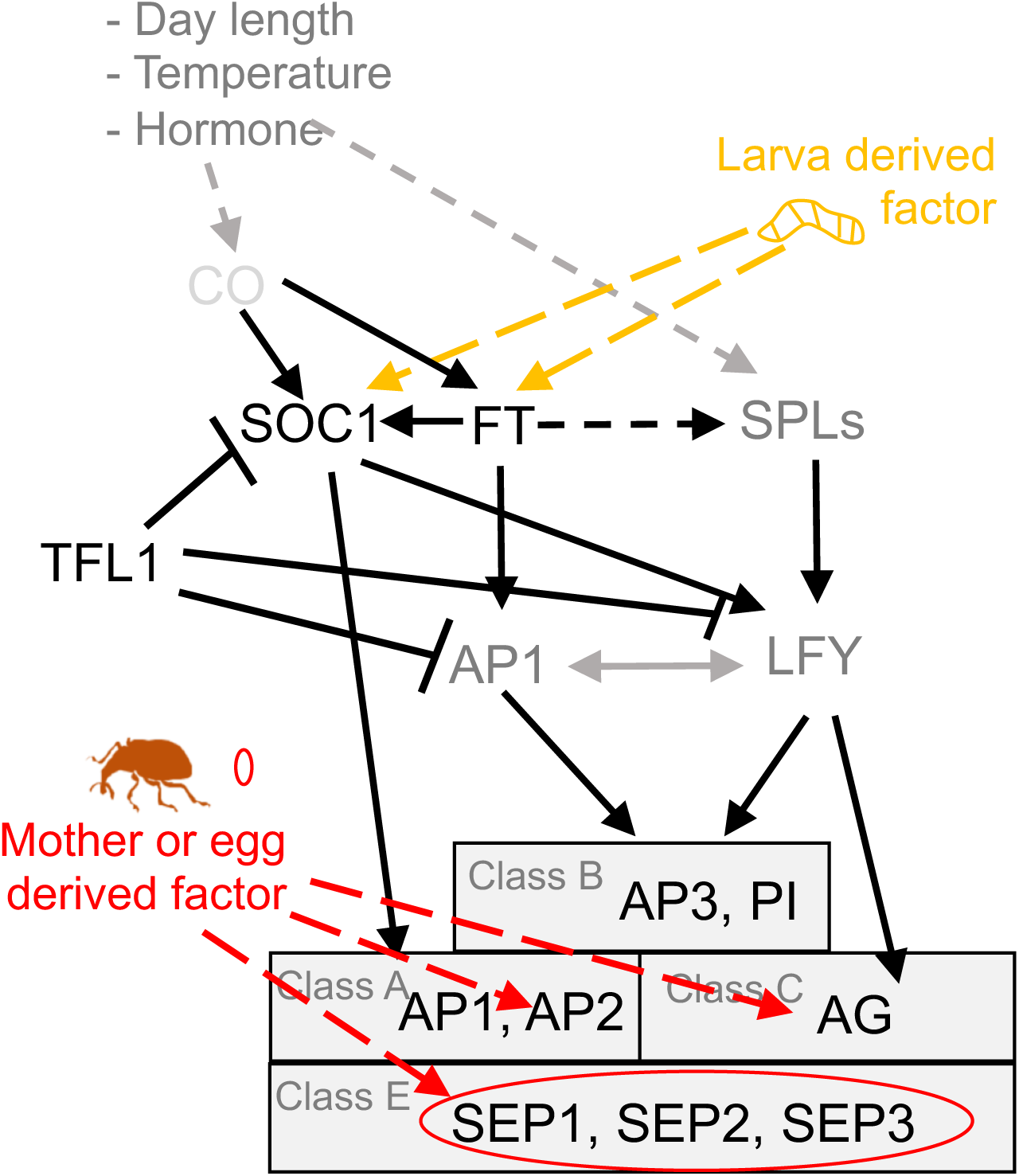
Flowering related signal pathway and interaction with *S. madaranus*. The flowering genes’ relationship was referred from Corbesier and Coupland (2006). Arrows and clasps indicate positive regulation and negative regulation of downstream genes respectively. Solid and dashed line indicate direct and indirect (or unknown) regulation.

**Supplemental Figure S6.**
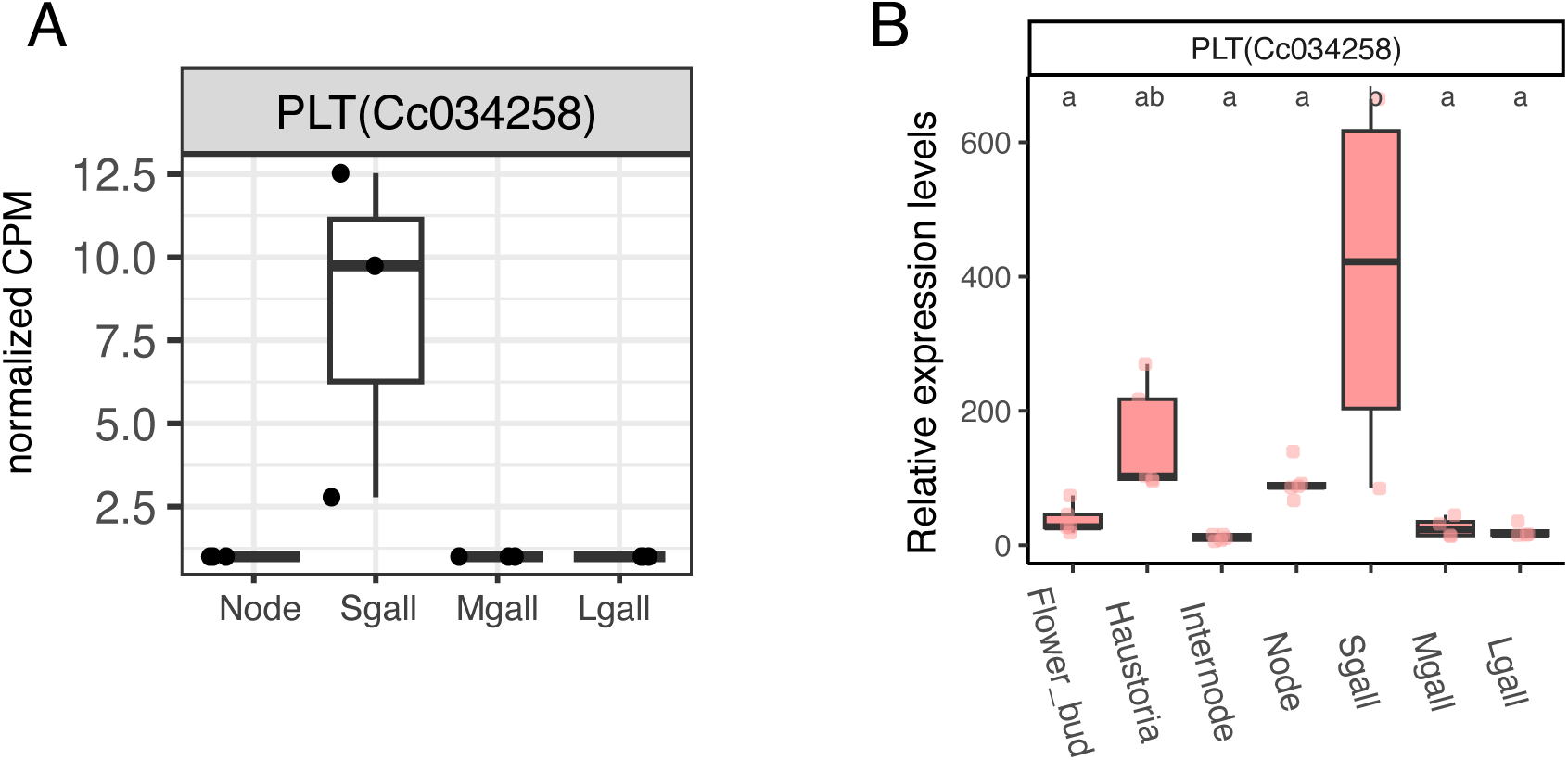
*CcPLT*-like gene expression pattern. (A) Expression levels in node before oviposition, Sgall, Mgall, and Lgall quantified as normalized CPM. (B) Expression levels in several tissues including galls quantified by qRT-PCR. Statistically significant differences are indicated by different letters above the boxes (ANOVA with Tukey’s HSD test, P < 0.05).

**Supplemental Figure S7.**
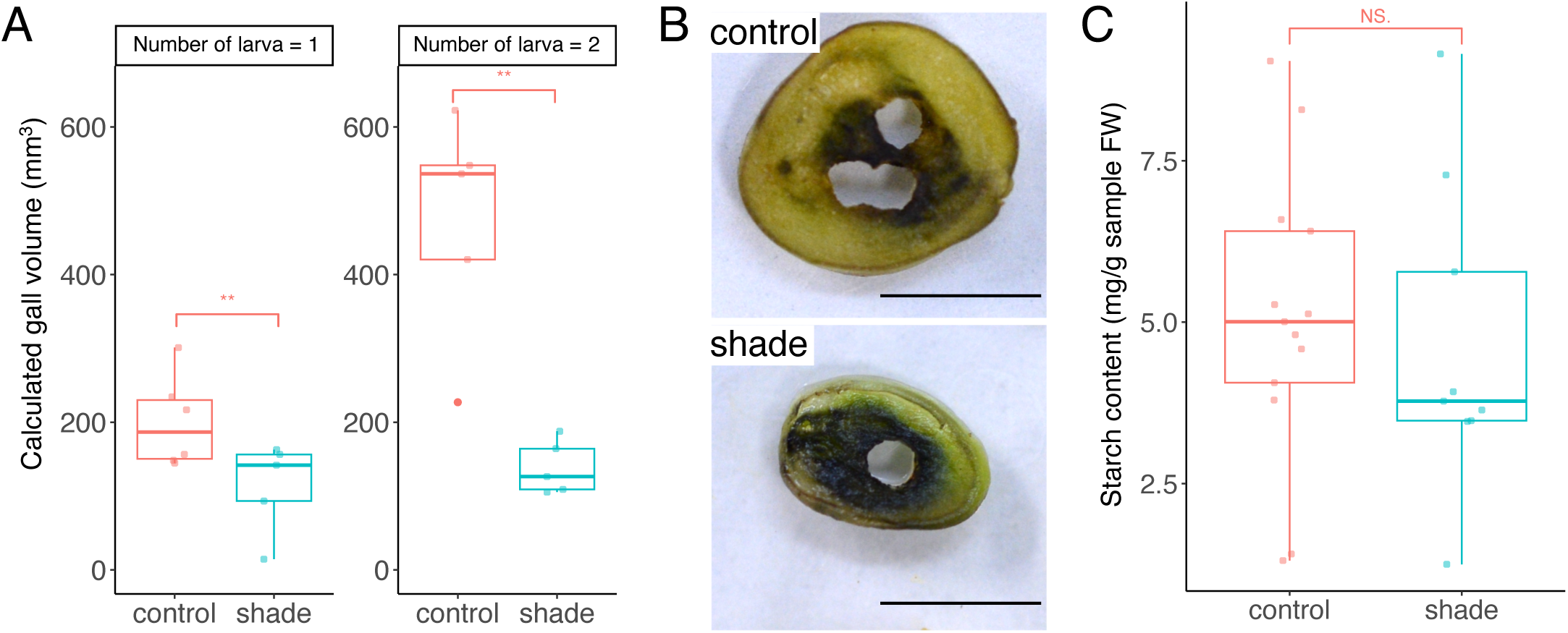
Influence of shading experiments on starch content. (A) Relationship between the gall size and number of larvae in the gall. (B) Starch in the galls stained blueblack with Lugol’s iodine solution. Scale bars, 5 mm. (B) Starch content quantification by using starch assay kit. **P < 0.01, NS, not significant difference; Wilcoxon rank sum test with Bonferroni correction

